# A neural moonshine conjecture. The representation of error-correcting codes in 3D symmetries invites the prospect of a Golay in the machine

**DOI:** 10.64898/2026.01.09.698680

**Authors:** Felipe Coelho Argolo

**Author notes:** F.A. contributed with idea inception, original draft and simulation. Author’s declare no competing interesting.

## Abstract

Conant-Ashby’s (‘good regulator’) theorem states that a simple regulator of a system must behave as an image of it. Notably, evolutionary features present in living beings often mirror natural processes, yielding symmetries between biological structures and the external environment. For instance, nervous cells can represent abstract entities through coordinated activities of neural networks. Central pattern generators (CPGs) used in locomotion display cyclic group symmetries (e.g. *Z_n_*), while the visual cortex and hyperbolic geometries in the hippocampus are respectively connected with Euclidean (*SE*(2), *SE*(3)) and Special linear (*SL*(2, *R*)) groups. This study renders conceivable a very efficient instance of the perfect error-correcting Golay code, whose automorphism group is the Mathieu sporadic simple group *M*_24_. We demonstrate that a set of 24 neurons is naturally organizable in a fully symmetric 3D polygonal network that embodies corruption-resistant dynamics. The existence of this configuration in biological systems is yet to be observed.

---

The mathematical field of Algebra uses symbolic expressions to represent the behavior of abstractions. Starting with polynomial equations for equalities between numerical quantities, it encompassed geometrical symmetries and arbitrary objects with the theory of groups (1). Sets of symbols and grammars with specific rules can mirror the corresponding abstract entities, enabling simple general approaches to understand underlying dynamics and examine specific scenarios. The application of algebraic formulations and group theory in natural sciences has been shown to be particularly effective (2), with physics frameworks being heavily based on the notion of symmetries and its formalization through groups, as in Noether’s theory and current formulations of the standard model of particle physics.

The study of symmetries is also at the basis of animal evolution and cognitive systems. Konrad Lorenz argued that the cognitive apparatus of an organism is itself an evolutionary adaptation to the reality of the physical world (3), while Paul Churchland suggested that the brain does not merely record data but constructs an internal compressed version of the external world’s abstract universals (4). Just as fins are shaped by the hydrodynamics of water, our neural structure would also reflect certain environmental dynamics. This biological structuralism aligns seamlessly with the Good Regulator theorem proposed by Conant and Ashby, which formally proves that any regulator that is maximally successful must be isomorphic to the system being regulated (5).

As expected, neural activity mimics the dynamics of several groups commonly used to model natural phenomena. Particularly, neural circuitry has shown to represent members of the “infinite groups”, such as cyclic and Euclidean groups. Conversely, mathematics also explores the symmetries of peculiar objects: the sporadic simple groups. At first, considered to be abstract curiosities, these entities were recently linked to practical applications.

This study demonstrates that the Mathieu group *M*_24_, a sporadic simple group of order ∼ 2 × 10^8^, can be efficiently embodied in a set of 24 spiking neurons. They spatially cover symmetrical vertices and faces of the great dodecahedron, and excitatory/inhibitory dynamics are based on local adjacency in this topology.

Due to the connection between *M*_24_, simple geometry and the perfect error-correcting binary Golay code, these structures may arise naturally and provide a basis for error correction in biological networks.

## Gait, space location and memory

*Z*_*n*_, *E*(*n*), *SL*(2, ℝ) The simplest biological rhythms, such as the gait of a centipede or the heartbeat of a leech, are governed by Central Pattern Generators (CPGs). These coupled oscillators display patterns described by cyclic groups (*Z*_*n*_) and dihedral groups (*D*_*n*_) (6, 7). For example, a network of neurons governing a tripod gait, is a physical instantiation of *Z*_3_ symmetry.

Pioneering work on grid cells and place cells in the hippocampus reveals a neural substrate for Euclidean groups (*E*(2) or *SE*(2)), allowing vector navigation and path integration (8). Recent theoretical work suggests that the geometry of these representations may be even more exotic, utilizing hyperbolic geometries, related to the Special Linear group *SL*(2, ℝ) (9).

## Moonshine, Sporadic Groups, and Biology

The classification of Finite Simple Groups was one of the monumental achievements of 20th-century mathematics. Among these, the “Sporadic” groups sit apart—exceptional structures that do not fit into the standard infinite families, they include Mathieu groups (*M*_11_, *M*_12_, *M*_22_, *M*_23_, *M*_24_), along with Janko, Conway, Fischer groups, the so called “Baby Monster” and “Monster”, known for their ridiculous order (M’s order is approximately 8 *×* 10^53^), and other groups. Originally considered to be pure mathematical abstractions, some sporadic simple groups were recently used to describe the fabric of reality: for instance, *M*_24_ group is linked to K3 surfaces, dimension-2 Calabi–Yau manifolds whose symmetries bridge different flavors of string theories in string duality. If sporadic groups describe deep symmetries in models of physical matter, could they also appear in biological systems?

This work suggests the physical existence of the Mathieu group *M*_24_ in neural cells. This comes from the fact that M24 is the automorphism group of the perfect error-correcting Golay code, which can conveniently be represented in 3D space through a symmetric polygon and local inhibitory/excitatory dynamics. An evolutionary pressure for efficient error correction would naturally converge upon the mathematical optimum defined by M24. Previous work has shown analog error-correcting models in grid cells of the entorhinal cortex (10). Similarly, the Golay code may represent a solution in the landscape of coding efficiency in embodied cognitive systems resistant to noise.

## The Golay Network Model

The Golay code is a (24, 12, 8) block code. It encodes 12 bits of data into a 24-bit word, capable of correcting any combination of up to 3 bit-flip errors. It is one of only two “perfect” binary codes (the other being the Hamming code).

It can be instantiated in a neural network consisting of 24 artificial neurons, indexed *i* ∈ {1, …, 24}, and adjacency defined by Golay generator matrix. Excitatory and inhibitory inter-neurons in this topology ensure the realization of the automorphic group of the Golay code, that is, M24. For the Golay code, the natural weight, number of active neurons, is often 8 (octads) or 12 (dodecad).

**Fig. 1.**
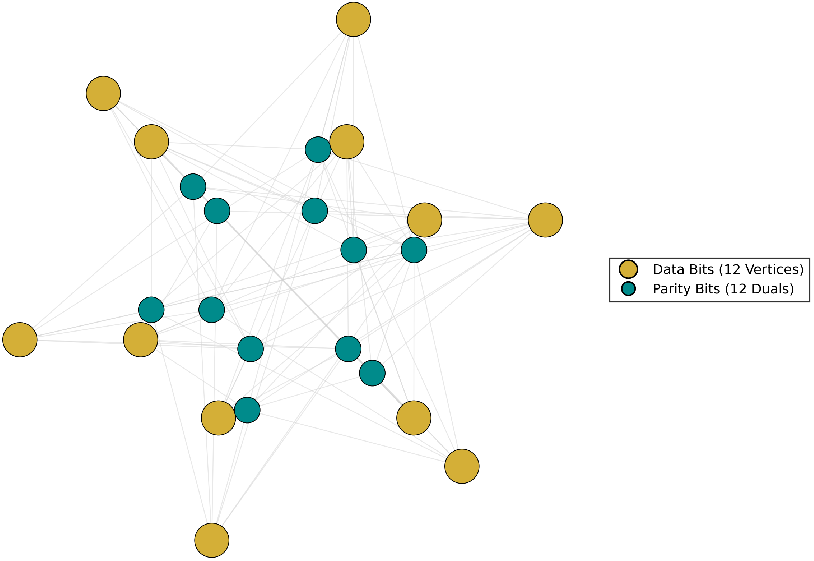
24 bits overlayed on the 3D geometry of the great dodecahedron

The great dodecahedron is a Kepler-Poinsot polyhedron that contains 12 edges and 12 pentagonal faces. It can be obtained by stellating the regular dodecahedron with pentagonal pyramids or by faceting the icosahedron. We may think of the network as possessing data neurons at the center of each pentagonal face and parity bits at the vertices. Whenever a data neuron from a pentagonal face is activated, it inhibits the 5 parity neurons located at the nearby vertices, activating the remaining 7 parity bits: one at the top at the pyramid and 6 opposite vertices in the antipodal side. Whenever a parity bit receives concomitant signals, they cancel out. This simple mechanism ensures that activating data neurons will produce valid Golay code-words.

## Discussion

The instantiation of M24 in a small set of neurons symmetrically displayed in a 3D polygon is a remarkable fact that increases the probability of its occurrence in nature.

Mathieu fist came across these groups when studying special symmetries of systems in which subsets of fixed size can be permuted. Inadvertently, he found the automorphism group of a perfect error-correcting code. These encodings optimize the number of parity bits, along with the possibility of locating errors and correcting them (reversion to original sequence of bits). Later on, the so-called Golay code was used for reliable information transmission in the Voyager space probes.

Unlike simple redundancy (repetition codes), which is metabolically expensive, the Golay code for error correction offers a “Goldilocks” solution to neural noise.

This connects back to the Good Regulator theorem. If the environment contains noise that follows specific statistical distributions, the optimal regulator must implement the optimal code to filter that noise.

Just like the hippocampus embodies hyperbolic geometry to pack memory, the motor cortex mimics cyclic groups to drive legs, and Euclidian groups represent space locations, neural circuits may also have converged to sporadic simple groups for information processing.

Future studies shall investigate the existence of such cell configurations. In addition, the hypothetical instantiation of other sporadic simple groups in biological structures remains to be outlined.

**Fig. 2.**
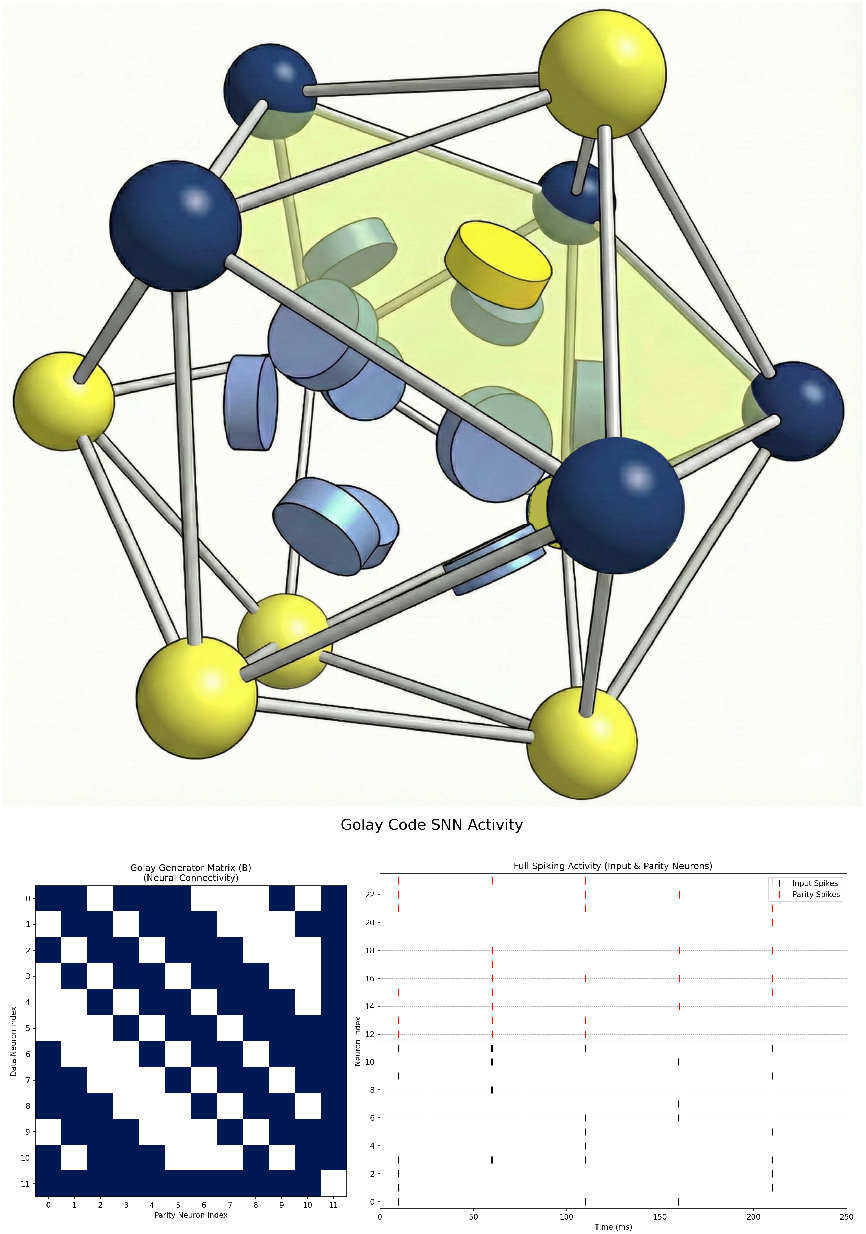
Top row displays an octad: an activated inner data bit and its 7 parity bits. One directly above the data bit and six in the antipodal side. Inspired by John Baez visualizations. Bottom-left panel shows the Golay Generator matrix, which corresponds to connectivity matrix of the network. The bottom-right panel shows inputs and corresponding parity bits configurations for several entries

## Materials and Methods

The encoder contains 24 neurons simulated with Brian 2 (v2.8.0.4 ; Python 3.10.12) for spiking neural networks (SNNs). It is a purely spike-driven, feed-forward network, with spiking times of 10 ms for activated unities. Its topology is defined by the Golay generator matrix. States of “parity neurons” are redefined whenever they are activated by a data neuron according to the rule: *x*_*t*_ = *x*_t−1_ + 1 mod 2. That is, whenever a new data neuron is activated, it propagates the signal to its parity neurons, performing a binary addition (in parallel) in the parity columns and ensuring that the final code-word (active neurons) are the active data neurons and the corresponding parity neurons after binary addition.

The decoder is able to parse 24-bit words, detect errors and correct them, although its architecture is more complex.

## Data, Materials, and Software Availability

All source code for the SNN simulation, data processing scripts, and visualization tools described in this paper are publicly available.

## Interactive network

The repository contains an interactive webpage in which one can activate arbitrary data bits and observe the corresponding parity bits in a great dodecahedral geometry. It also generates valid code-words and features a corruption injection and error-correction procedure.

## Automorphic group demonstration

The repository contains an animated random-walk that performs arbitrary actions from M24, showing the automorphic group of the Golay code.

The full repository, including detailed documentation and instructions to reproduce all figures and experiments, can be accessed at:

https://github.com/fargolo/neural-moonshine

The code is released under the MIT License.

## ACKNOWLEDGMENTS

Financial support - FAPESP- PRIP. John Baez for inspiring visualizations https://blogs.ams.org/visualinsight/2015/12/01/golay-code/

